# Rapid clearing of biological organs by using phosphoric acid, a hydrophilic solution with high refractive index

**DOI:** 10.1101/452425

**Authors:** Masakazu Umezawa, Shinsuke Haruguchi, Rihito Fukushima, Shota Sekiyama, Masao Kamimura, Kohei Soga

## Abstract

Tissue clearing is a fundamental challenge in biology and medicine to achieve high-resolution optical imaging of tissues deep inside intact organs. The clearing methods, reported up to now, require long incubation time or physical/electrical pressure to achieve tissue clearing, which is done by matching the refractive indices of the whole sample and medium to that of the lipid layer. Here we show that phosphoric acid increases the refractive index of the medium and can increase the transparency of formalin-fixed tissue samples rapidly. Immersion of fixed tissues of mice in phosphoric acid solutions increased their transparency within 60 min in the case of 3-mm-thick fixed tissue specimens. While phosphoric acid suppresses bright signals on the boundary of cells in their phase-contrast images, it does not damage the morphology of cell membrane with phospholipid bilayer. The protocol presented herein may contribute to develop better and faster soaking methods for tissue clearing than previously reported protocols.

**Highlights:**

▪ Phosphoric acid can reduce light scattering by tissue samples.
▪ Tissue clearing effect of phosphoric acid is fast and needs only 60-min incubation.
▪ Cell membrane was preserved during incubation using phosphoric acid.

## Introduction

Whole-tissue and whole-body imaging of single cells in opaque organisms like mammals has been fundamental challenges in biology and medicine. Biological three-dimensional imaging has been achieved by reconstructing three-dimensional structure from images taken from mechanically serially sectioned tissues, for example serial block-face scanning electron microscopy [1] and array tomography [2]. However, the technique is not only labor-intensive and time-consuming but also error-prone [3]. The utility of rendering tissue optically transparent has been well known for three-dimensional imaging of cell populations located deep inside intact tissue. Optical sectioning provides a potentially fast, simple and inexpensive alternative for three-dimensional reconstruction; however, its utility for deep imaging is prevented by tissue opacity.

Optical imaging of thick tissues is mostly limited by scattering of imaging light through thick tissues, which contain various cellular and extracellular structures with different refractive indices. The imaging light traveling through different structures scatters and loses its excitation and emission efficiency, resulting in a lower resolution and imaging depth [4]. The low refractive index of water compared with that of cellular structures containing proteins and lipids has been the major issue in this regard; in order to address this problem, the main concept of tissue clearing is to adjust the difference between the refractive index of the medium and that of the lipid bilayer [5, 6]. The clearing was achieved by hydrophobic solutions during the early stage of technical development [7]. In this method, a fixed sample is incubated in a mixture of benzyl-alcohol and benzyl benzoate after dehydration of samples with ethanol and hexane. Subsequent reports have shown that the use of tetrahydrofuran (THF) or dibenzyl ether (DBE) instead of alcohol for dehydration improved the clearing of entire brains of adult mice [8, 9] and this method is applicable to clearing lipid-rich spinal cord [10]. Removing membrane lipids by repetitive organic solvent-based dehydration-rehydration improved the permeabilization of tissues [11].

To reduce the denaturation and to improve the retention of fluorescent signals in formalin-fixed tissue samples, urea-based hydrophilic mixtures, Sca*l*e, have been developed [12]. Because this technique provided a simple, inexpensive alternative to array tomography and serial section electron microscopy, it contributed to promoting and devising further protocols for tissue clearing. Formamide or formamide/poly(ethylene glycol), Clear^T^ and Clear^T2^, were able to clear mouse embryo and brain with no detergents or solvents [13]. SeeDB containing optical clearing agents, fructose and α-thioglycerol, renders tissue samples transparent to allow analyses of cellular morphology without removing any components of tissues during the clearing process and thus with minimum deformation artifacts [14]. Adding fructose to urea can control ureamediated tissue expansion during the clearing process [15]. Sugar and sugar-alcohol solutions also achieve clearing of 100-μm-thick mouse brain slices [16]. Preparation of clearing solutions to adjust the refractive index of samples to that of immersion oil (1.518) on the objective lens is effective for high-resolution fluorescence imaging [17]. Electrophoresis to remove lipids after embedding tissue into hydrogel polymer contributes to better tissue-clearing in a method named CLARITY [18]. However, because this method was technically complex, a passive tissue clearing approach using different fixative-hydrogel monomer crosslinking solutions was subsequently reported for adult rodents or human postmortem brain [19, 20]. Solution of 2,2’- thiodiethanol is also effective for clearing brains [21] and facilitated three-dimensional observation of formalin-fixed and fluorescent-labeled whole mouse brains and 2-mm-thick block of human brains in combination with CLARITY [22]. To improve the methodology to deliver reagents deep inside thick tissues, rapid (4-20 h for clearing whole organs) technique of active clarity, ACT-PRESTO, was reported for rendering large tissue samples such as rat and rabbit brains, and even whole bodies of adult mice, optically transparent [23]. On the other hand, CUBIC method achieves tissue transparency by passively clearing phospholipids and is compatible with hydrogel embedding using a mixture of urea and aminoalcohol [24], which decolorizes blood by efficiently eluting the heme chromophore from hemoglobin [25]. Optimized CUBIC protocols for whole-body transparent observation of mice [26] and organs of rats [27] are also reported. To reduce tissue damage and degradation, Sca*l*eS method was developed with a mild tissue-permeant sugar-alcohol, sorbitol, and allowed optical reconstructions of three-dimensional mapping of amyloid plaques, neurons and microglia by successive fluorescence and electron microscopy [28]. However, one of the major issues is that the protocols reported above for clearing tissues require long incubation time from several hours to days or pressure that also damages the samples. Rapid and simple soaking protocols for tissue clearing are thus desirable.

Many hydrophilic solvents for tissue clearing, Sca*l*e, FRUIT, and CUBIC, contain liquids of high refractive index to increase the refractive index of the incubation medium, in some cases, surfactant to delipidate [29] and potentially promote infiltration of the reagent deeper into tissues, and urea [12, 15, 24, 28, 30]. The refractive index of a material is proportional to the square root of the dielectric constant of that material. Therefore, we hypothesized that tissue clearing is achieved by not only increasing the refractive index of the medium but also by affecting polarization and dielectric coefficient of the lipid membrane, which is composed of non-polarized fatty-acid chain and polarized phosphate group, due to some molecules such as urea in the solutions. In particular, the research group who developed Sca*l*e containing urea, Triton X-100, and glycerin, showed that partial tissue clearing was achieved using only urea and the surfactant [12]. The mechanism underlying tissue clearing effect by urea is not known but may be important to explore. In the present study, we first investigated the potential contribution of urea to adjusting the refractive indices of the medium and lipid membrane. Next, the tissue clearing effect of phosphoric acid, an anionic molecule that does not interact with anionic lipid bilayer but increases the refractive index of medium, was investigated. We show that urea does not directly contribute to decreasing the refractive index of lipid bilayer, and phosphoric acid, which increases the refractive index of the medium, is able to clear the tissues. Although the incubation for fixed tissue sample in phosphoric acid is only passive immersion, the incubation time required for clearing whole organ samples of mice is faster (≈ 1 h) compared to the case of other clearing reagents. The rapid optical clearing protocol using phosphoric acid will contribute to further improvement of tissue clearing protocols and will aid in the advancement of biological research.

## Materials and Methods

### Measurement of refractive index of solutions

Solutions of saturated concentration of 1,2-dipalmitoyl-sn-glycero-3-phosphocholine (DPPC) and 4 M urea were prepared in ethanol. The refractive indices of the mixed solutions in ethanol, water, and aqueous solutions of phosphoric acid (4–14.2 M), urea (4 M), sodium iodide (2 and 4 M), indium chloride (4 M), and ScaleA2 were measured by Abbe’s refractometer (NAR-1T, Atago Co., Ltd., Tokyo, Japan).

### Measurement of refractive index and optical loss of liposome suspension

Liposome suspension was prepared by the Bangham method [31]. Briefly, a mixture of 12 pmol of DPPC, 12 μmol of 1,2-dioleoyl-sn-glycero-3-phosphocholine (DOPC), 12 μmol of cholesterol, and 4 μmol of *N*-(Carbonyl-methoxypolyethyleneglycol 2000)-1,2-distearoyl-sn-glycero-3-phosphoethanolamine (DSPE-PEG) was prepared in 1600 μL of chloroform in an eggplant-shaped glass flask, and then evaporated at 37°C to remove chloroform and to obtain a thin lipid bilayer on the inner wall of the flask. 7.5-mL of either of distilled water, aqueous solution of urea (4 M) or phosphoric acid (14.2 M) was added into the flask and sonicated to prepare liposome suspension in each liquid. The light transparence (wavelength: 600 nm) of the liposome suspensions were measured by spectrometer V-660 (JASCO Co., Tokyo, Japan) and 1.5-mL disposable cuvette (Brand GmbH & Co. KG, Wertheim, Germany).

### Measurement of optical loss of cell suspension of mouse brain

All animals were handled in accordance with the national guidelines for the care and use of laboratory animals and with the approval of the Animal Care and Use Committee of the Tokyo University of Science. Brain samples isolated from adult ICR mouse sacrificed under hyperanesthesia were minced by using BioMasher (Nippi Inc., Tokyo, Japan), suspended in PBS, and fixed by 4 wt% formaldehyde in PBS overnight. Fixed cell suspension was collected by centrifugation (1200 ×g, 5 min) and washed with PBS three times, and divided into different tubes. The cell suspensions were then incubated in either of Sca*l*eA2, PBS, or aqueous solution of urea (4 M), indium chloride (4 M), sodium iodide (2, 4 M), Triton X-100 (10 wt%), phosphoric acid (8.5, 11.4, 14.2 M), or 0.1 wt% Triton X-100 in phosphoric acid (14.2 M) for 24 h at room temperature. The optical loss (OD600) was measured by spectrometer V-660 (JASCO) and 1.5-mL disposable cuvette (Brand GmbH & Co. KG).

### Observation of morphology and phase-contrast image of formalin-fixed cells

Cultured Colon-26 cells were maintained in Minimum Essential Medium [(+)Eagle’s salt, (+)L-glutamine] (Gibco, Thermo Scientific, MA, USA) supplemented with 10% fetal bovine serum (Biowest, Nuaillé, France) and 1% penicillin-streptomycin (Wako, Tokyo, Japan) at 37°C in humidified air containing 5% CO_2_. After staining of the cell membrane by CellMask Orange (Thermo Fisher Scientific), the cells were fixed by 4 wt% formaldehyde in PBS for 10 min, washed with PBS, and then incubated in phosphoric acid (14.2 M) for 0-48 h. Cell membrane stained with CellMask was observed by fluorescence microscope through a band-pass filter (cut-on wavelength: 575 nm) under excitation (540 ±10 nm). The cells without staining were also fixed with formalin, washed with PBS, incubated in ScaleA2, phosphoric acid (14.2 M), or PBS for 24 h, and observed under phase-contrast microscopy.

### Measurement of optical loss of mouse tissues

Brain, liver, kidney, and lung samples were obtained from adult ICR mouse under anesthesia. The samples of liver, kidney, and lung were fixed by 4 wt% formaldehyde in PBS for 48 h, washed with PBS, and cut into approximately 3-mm-thick specimens. The fixed tissue specimens and brain hemispheres of mice were incubated in either of ScaleA2, PBS, or aqueous solution of indium chloride (4 M), sodium iodide (4 M), Triton X-100 (10 wt%), or phosphoric acid (8.5, 11.4, 14.2 M) for 60 min. The macroscopic images of the treated samples were captured by a digital camera. To confirm the reversibility of tissue clearing, 450-pm-thick specimens of mouse brain were prepared, fixed by formaldehyde for 60 min, washed with PBS, incubated in 14.2 M phosphoric acid or PBS for 60 min, replaced and kept in PBS for 24 h, and their images were also recorded by digital camera.

## Results and Discussion

Phospholipid containing both hydrophobic and hydrophilic parts is arranged in two layers on the plasma membrane; the hydrophobic part is on the interior of the membrane, whereas the hydrophilic part points outwards, toward either the cytoplasm or the fluid that surrounds the cell. The hydrophobic and hydrophilic parts have different refractive indices due to the difference in their dielectric constant. Our hypothesis was a possible contribution of decrease in polarization of the phosphate group, induced by electrostatic interaction of the polarized group with urea, to decrease the refractive index difference between the lipid bilayer and medium. The refractive index of the solution of a representative phopholipic dipalmitoylphosphatidylcholine (DPPC) in ethanol, which is able to dissolve both DPPC and urea, was higher than those of ethanol and water (Fig 1a). Adding urea to ethanol and DPPC solution in ethanol increased their refractive indices (Fig 1a). These results suggest that urea does not decrease the refractive index of DPPC in ethanol but increases the refractive indices of the solutions. The light transparency of liposome suspended in aqueous solution of urea (4 M), which shows higher refractive index than water, was higher than that of liposome suspended in water (Fig 1b). Moreover, 14.2 M phosphoric acid, which is an anionic molecule that does not interact with the anionic phosphate group of lipid bilayer, shows higher refractive index than 4 M urea solution and increased transparency of the liposome (Fig 1b). We concluded that urea increased the transparency of liposome not by affecting the polarization or refractive index of phosphate group of the lipid bilayer, but by reducing the difference in refractive indices between the medium and the lipid bilayer as well as phosphoric acid. The clearing effect of phosphoric acid is greater than that of 4 M urea due to higher refractive index.

**Fig 1.**
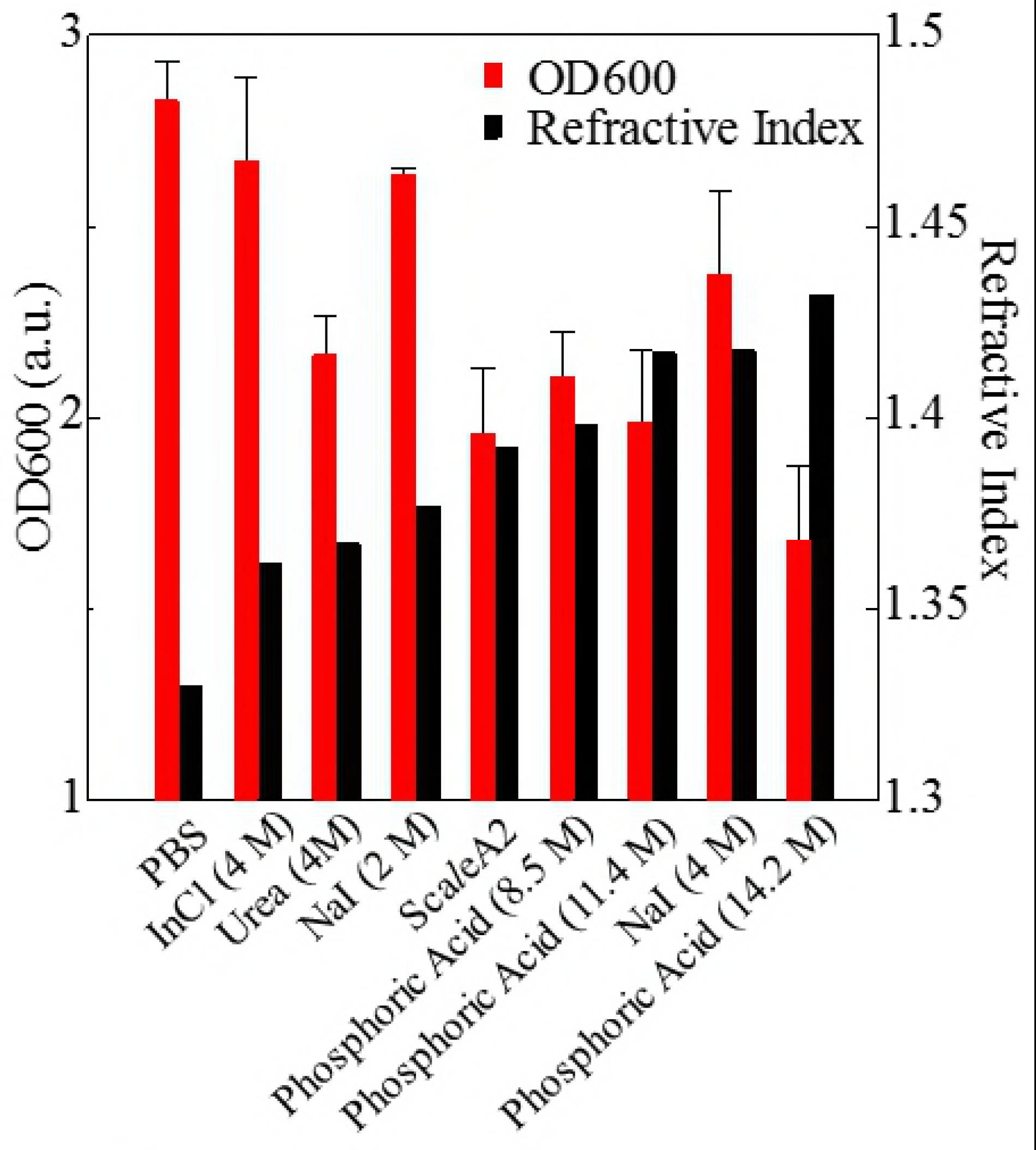
Effects of urea and phosphoric acid on the refractive indices of solvents and on transparency of liposome suspension. Refractive indices of DPPC in ethanol and its mixture with urea are shown in (a). An increase in the refractive index of DPPC solution in ethanol by urea suggests that urea does not influence the polarization of the phosphoric acid group of lipid bilayer but increases the refractive index of the medium. The effects of urea (4 M) and phosphoric acid (14.2 M) on the refractive index of the medium and transparency of liposome, determined by OD600, are shown in (b). The data of transmittance (%) are shown as mean ±SD (n = 3). Abbreviation: DPPC, 1,2-dipalmitoyl-sn-glycero-3-phosphocholine.

Next, we investigated the clearing effect of phosphoric acid, which decreased the light loss of the liposomes, and other candidate chemical solutions on fixed cell suspensions of mouse brain tissues. We targeted phosphoric acid solution, because phosphate group composes of the strongly polarized part and thus contributes to high refractive index of the phospholipid bilayer. Tissue-clearing effects of many kinds of solvents were evaluated by measuring the light loss (OD600) of suspension of tissue cells fixed with formaldehyde after homogenization, as shown in previous studies [24]. The light loss of formalin-fixed cell suspensions of mouse brain was decreased by 24-h incubation in phosphoric acid and Sca*l*eA2 (Fig 2). On the other hand, although 4 M sodium iodide (4 M) solution has a similar high refractive index to phosphoric acid, sodium iodide did not decrease the light loss of the cell suspension during the 24-h incubation. Indium chloride (4 M) also has a similar refractive index to urea (4 M), it did not attenuate the light loss by the 24-h incubation as well as sodium iodide. This lack of clearing effect of sodium iodide and indium chloride may be due to relatively poor infiltration of this molecule into fixed tissues or cells. Incubation of cultured cells in tissue clearing solutions (phosphoric acid and Sca*l*eA2) attenuated the bright signal on the boundary of cells in the phase-contrast image in a concentration-dependent manner (Fig 3). Because the bright signal on the cell boundary is generated where the difference in refractive index between the sample and background is large, the decrease in the signal by the clearing solutions confirms the effects of the solution on the refractive index difference between the cell membrane and medium. The stability of formalin-fixed cell membrane in phosphoric acid was confirmed by labeling the membrane of the cultured cells, suggesting that phosphoric acid does not damage the fixed membrane during incubation (Fig 4). Macroscopic images of formalin-fixed organ specimens (liver, kidney, and lung) of mice demonstrated that the samples were partially cleared by 60min incubation in phosphoric acid solution (Fig 5). Phosphoric acid (8.5 M) is effective for reducing scattering by the liver, while higher concentrations of 11.4 M and 14.2 M are needed for the kidney and lung, respectively. On the other hand, 60-min incubation was too short to clear the tissue specimens by Sca*l*eA2. Rapid tissue clearing by phosphoric acid is possibly due to fast infiltration of the molecule because of its small size. The molecular size (volume) of phosphoric acid (PO_4_ ^3−^) is smaller than that of urea (CH_3_COCH_3_), glycerol [CH_2_(OH)- CH(OH)-CH_2_(OH)], and sugars and sugar alcohols. Other candidate chemicals investigated in the present study, sodium iodide (4 M) and indium chloride (4 M), did not make the tissues transparent but induced shrinkage of samples, especially the kidney specimen. Phosphoric acid is also able to make brain hemisphere transparent as shown in Fig 6. The clearing by phosphoric acid is reversible; re-incubation of transparent 450-μm-thick brain slices in PBS made the sample opaque as well as the original image before phosphoric-acid treatment (Fig 6b). This result indicates that phosphoric acid reduces light scattering in the tissue sample not by irreversible chemical modification but by adjusting the refractive index of the sample to a value close to that of the cell membrane by simple infiltration. Re-incubation of the sample in PBS removed phosphoric acid from the sample and made it opaque reversibly.

**Fig 2.**
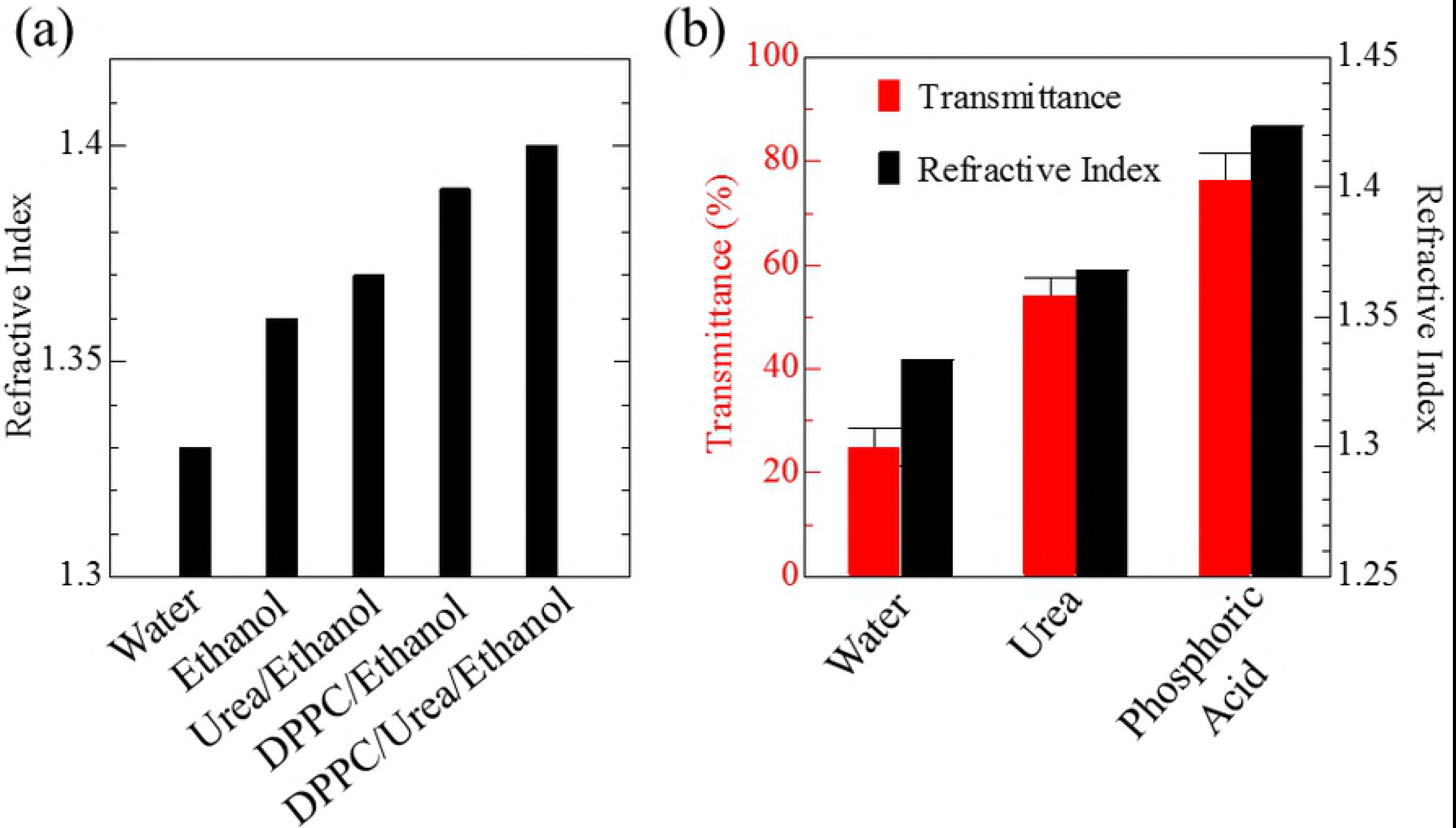
Effects of candidate solutions with different refractive indices on light loss for fixed cell suspension. Effects of 24-h incubation in candidate chemical solutions with various refractive indices on OD600 values (mean ±SD, n = 3) of brain cell suspension fixed with formaldehyde are shown. Phosphoric acid (8.5, 11.4, 14.2 M) further increases the refractive indices of medium and transparency of the cell suspension than sodium iodide (4 M), urea (4 M), and indium chloride (4 M). Abbreviations: InCl, indium chloride; NaI, sodium iodide.

**Fig 3.**
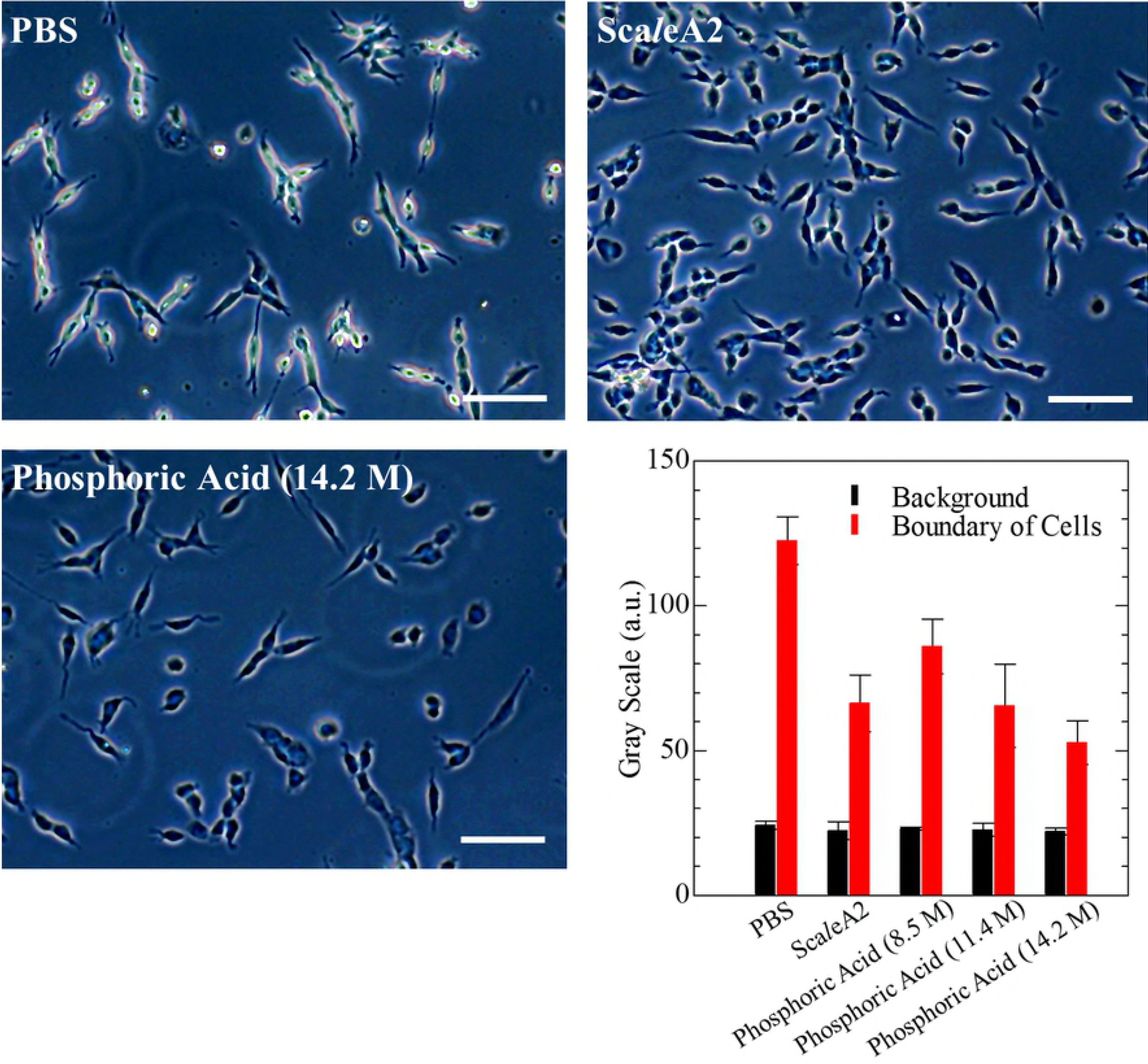
Phase-contrast images of formalin-fixed cultured Colon-26 cells incubated in phosphoric acid or Sca*l*eA2. Effect of Sca*l*eA2 and phosphoric acid (14.2 M) solution on the phase contrast on the edge of cultured cells is shown in the microscopic images. Scale bars indicate 50 pm. The graph shows the concentration-dependent effect of phosphoric acid (8.5-14.2 M) on the gray values of background and the edge in the microscopic images.

**Fig 4.**
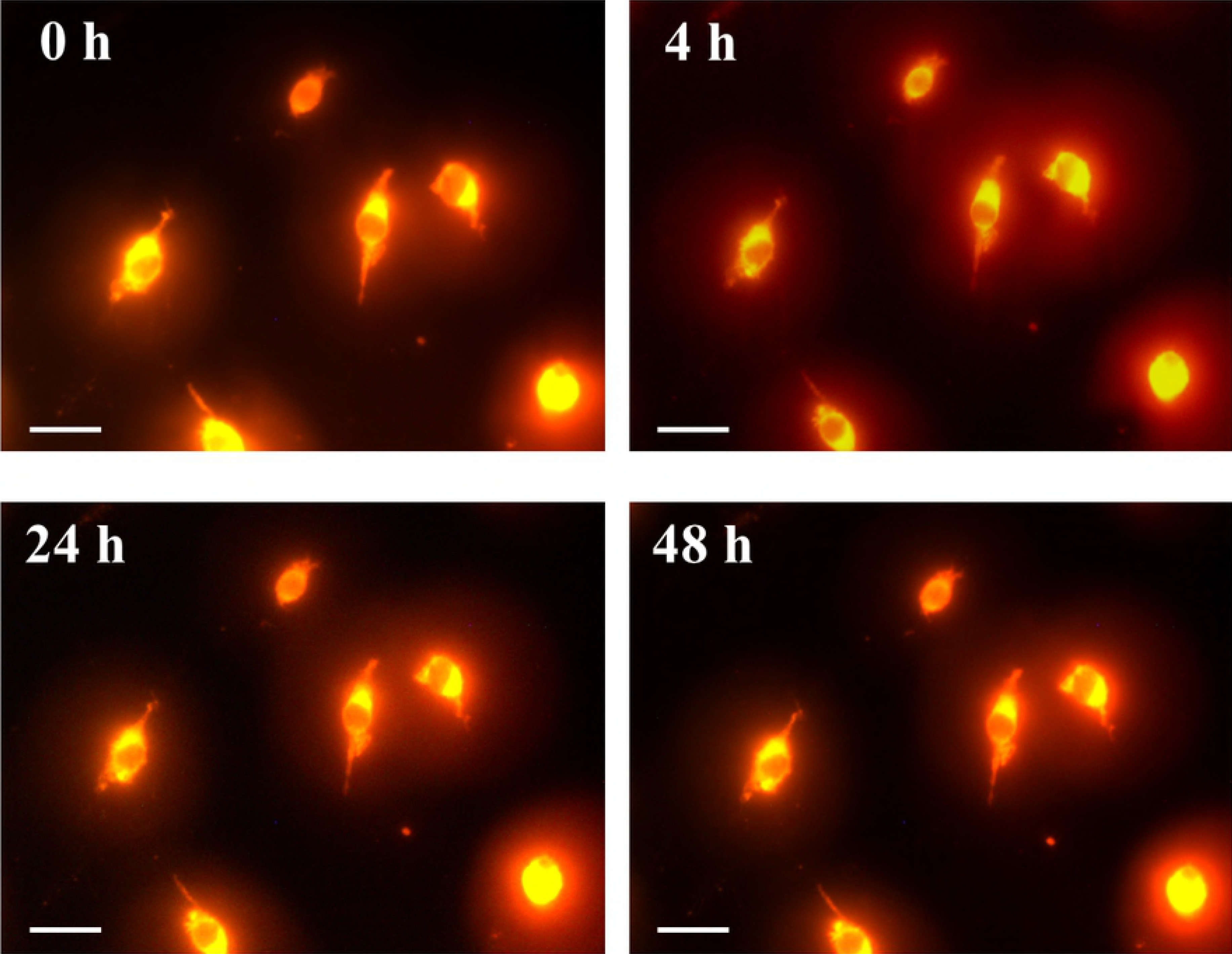
Images of formalin-fixed plasma membrane of cultured cells incubated in phosphoric acid. Cultured Colon-26 cells were stained with CellMask Orange (Thermo Fisher Scientific), fixed by 4 wt% formaldehyde in PBS, washed with PBS, and then incubated in phosphoric acid (14.2 M) for 0-48 h and observed under microscope. Scale bars indicate 10 pm. The images confirmed that formalin-fixed cell membrane was retained during incubation in the solution.

**Fig 5.**
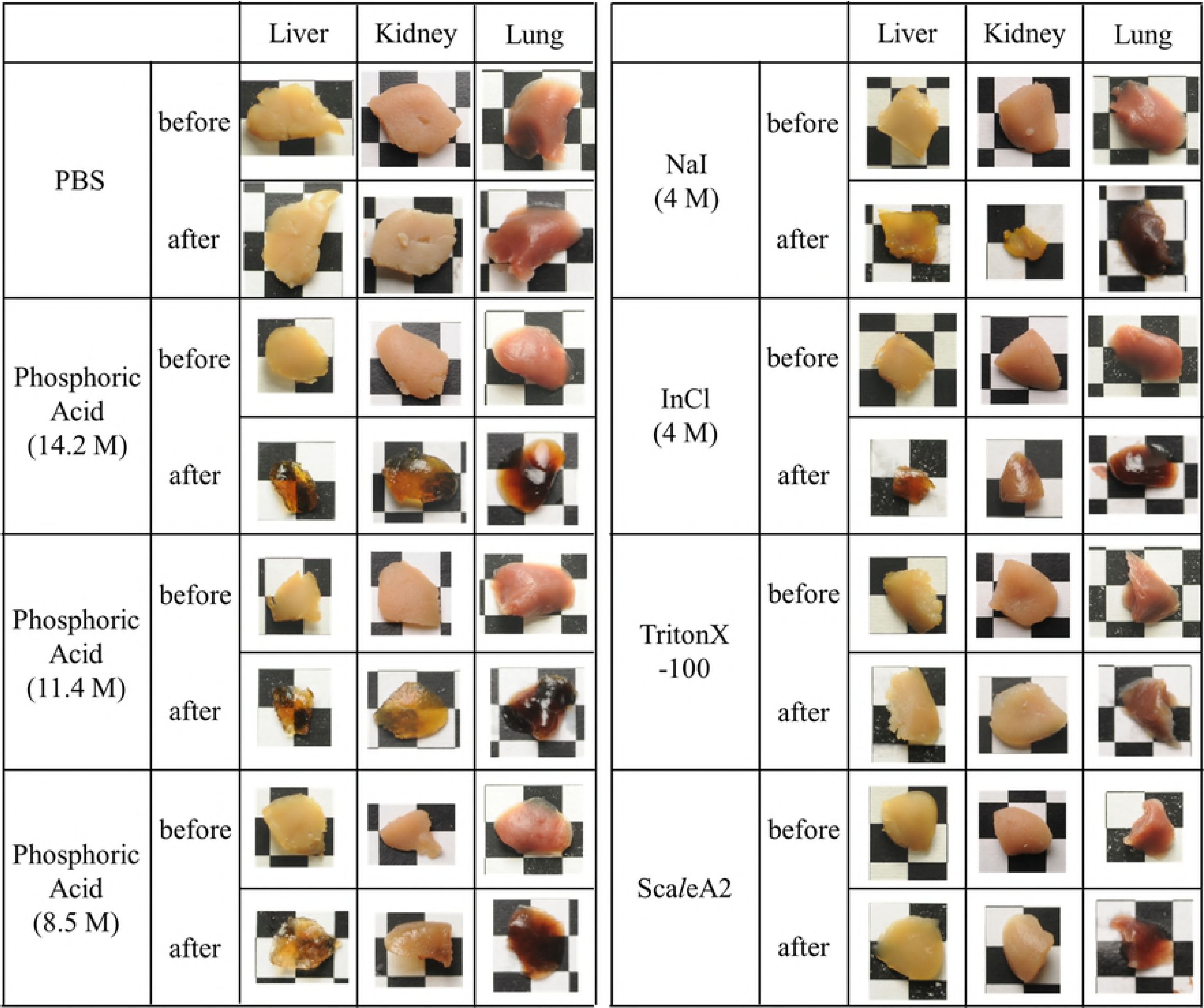
Effect of incubation in chemical solutions on macroscopic images of formalin-fixed tissue specimens of mice. The organ specimens of murine (a) liver, (b) kidney, and (c) lung were fixed with 4 wt% formaldehyde for 48 h, washed with PBS, and then incubated in indicated chemical solutions for 60 min. The apertures of the background lattice are 4 mm.

**Fig 6.**
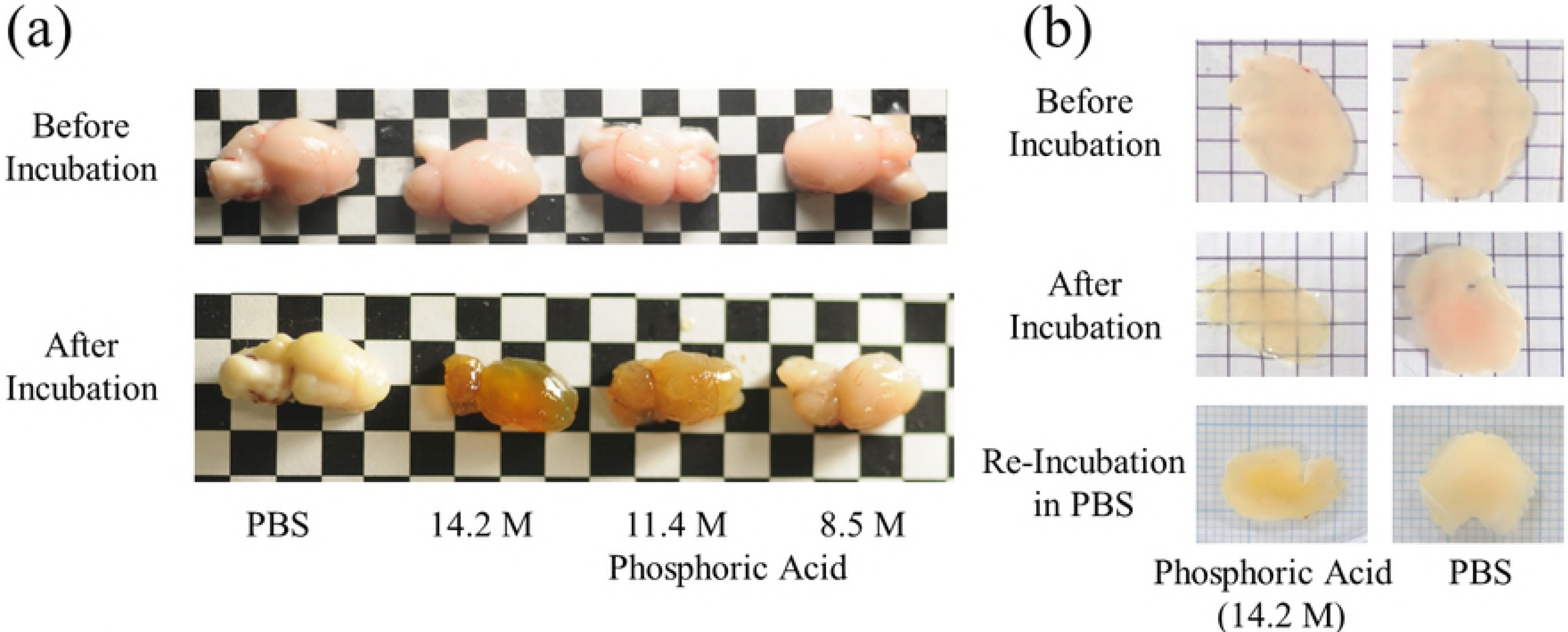
Clearing mouse brains by phosphoric acid. Representative images of (a) hemispheres and (b) 450-pm-thick specimens of mouse brains before and after 60-min incubation in phosphoric acid (8.5, 11.4, and 14.2 M) are shown. As shown in (b), the brain tissue that were cleared once by phosphoric acid (14.2 M) returned to opaque state by re-incubation in PBS for 24 h.

## Conclusions

The present study showed that phosphoric acid, a hydrophilic solvent, can clear tissues including brain, liver, kidney, and lung of mice. Phosphoric acid reduces light scattering by mouse tissues rapidly as it requires only 60-min incubation, while it does not damage the morphology of cell membrane with phospholipid bilayer. The present study showed that phosphoric acid reduced scattering by mouse tissues but did not reduce the light loss due to absorption by the tissues. Combination use of decolorization reagent such as aminoalcohol [25] with small molecular solutes that increase refractive index, like phosphoric acid, may achieve better and rapid tissue clearing. The potential of phosphoric salts with neutral pH for tissue clearing is also of future interest because acidic condition generally denatures fluorescent proteins [32, 33]. The rapid reducing of scattering with the use of phosphoric acid as presented here will contribute to develop better and faster soaking methods for tissue clearing that previously reported protocols.

## Supporting Information

The data required to reproduce the optical loss of suspensions of liposome and brain cells in each solvent are posted as a supporting information.

## References

[1] Denk W, Horstmann H. Serial block-face scanning electron microscopy to reconstruct threedimensional tissue nanostructure, PLoS Biol. 2004;2: e329.

[2] Micheva KD, Smith SJ. Array tomography: a new tool for imaging the molecular architecture and ultrastructure of neural circuits. Neuron. 2007;55: 25–36.

[3] Helmstaedter M, Briggman KL, Denk W. 3D structural imaging of the brain with photons and electrons. Curr Opin Neurobiol. 2008;18: 633–641.

[4] Boas D. A fundamental limitation of linearized algorithms for diffuse optical tomography. Opt Express. 1997;1: 404–413.

[5] Genina EA, Bashkatov AN, Tuchin VV. Tissue optical immersion clearing. Expert Rev Med Devic. 2010;7: 825–842.

[6] Richardson DS, Lichtman JW. Clarifying Tissue Clearing. Cell 2015;162: 246–257.

[7] Dodt HU, Leischner U, Schierloh A, Jährling N, Mauch CP, Deininger K, et al. Ultramicroscopy: three-dimensional visualization of neuronal networks in the whole mouse brain. Nat Methods. 2007;4: 331–336.

[8] Becker K, Jährling N, Saghafi S, Weiler R, Dodt HU. Chemical clearing and dehydration of GFP expressing mouse brains. PLoS One. 2012;7: e33916.

[9] Ertürk A, Mauch CP, Hellal F, Förstner F, Keck T, Becker K, et al. Three-dimensional imaging of the unsectioned adult spinal cord to assess axon regeneration and glial responses after injury. Nat Med. 2012b;18: 166–171.

[10] Ertürk A, Becker K, Jährling N, Mauch CP, Hojer CD, Egen JG, et al., Three-dimensional imaging of solvent-cleared organs using 3DISCO. Nat Protoc. 2012a;7: 1983–1995.

[11] Renier N, Wu Z, Simon DJ, Yang J, Ariel P, Tessier-Lavigne M. iDISCO: a simple, rapid method to immunolabel large tissue samples for volume imaging. Cell. 2014;159: 896–910.

[12] Hama H, Kurokawa H, Kawano H, Ando R, Shimogori T, Noda H, et al. Scale: a chemical approach for fluorescence imaging and reconstruction of transparent mouse brain. Nat Neurosci. 2011;14: 1481–1488.

[13] Kuwajima T, Sitko AA, Bhansali P, Jurgens C, Guido W, Mason C. ClearT: a detergent- and solvent-free clearing method for neuronal and non-neuronal tissue. Development. 2013;140: 1364–1368.

[14] Ke MT, Fujimoto S, Imai T. SeeDB: a simple and morphology-preserving optical clearing agent for neuronal circuit reconstruction. Nat Neurosci. 2013;16: 1154–1161.

[15] Hou B, Zhang D, Zhao S, Wei M, Yang Z, Wang S, et al. Scalable and DiI-compatible optical clearance of the mammalian brain. Front Neuroanat. 2015;9: 19.

[16] Yu T, Qi Y, Wang J, Feng W, Xu J, Zhu J. Rapid and prodium iodide-compatible optical clearing method for brain tissue based on sugar/sugar-alcohol. J Biomed Opt. 2016;21: 081203.

[17] Ke MT, Nakai Y, Fujimoto S, Takayama R, Yoshida S, Kitajima TS, et al. SuperResolution Mapping of Neuronal Circuitry With an Index-Optimized Clearing Agent. Cell Rep. 2016;14: 2718–2732.

[18] Chung K, Wallace J, Kim SY, Kalyanasundaram S, Andalman AS, Davidson TJ, et al. Structural and molecular interrogation of intact biological systems. Nature. 2013;497: 332–337.

[19] Tomer R, Ye L, Hsueh B, Deisseroth K. Advanced CLARITY for rapid and high-resolution imaging of intact tissues. Nat Protocol. 9: 2014;1682–1697.

[20] Zheng H, Rinaman L. Simplified CLARITY for visualizing immunofluorescence labeling in the developing rat brain. Brain Struct Funct. 2016;221: 2375–2383.

[21] Aoyagi Y, Kawakami R, Osanai H, Hibi T, Nemoto T. A rapid optical clearing protocol using 2,2’-thiodiethanol for microscopic observation of fixed mouse brain. PLoS One. 2015;10: e0116280.

[22] Costantini I, Ghobril JP, Di Giovanna AP, Allegra Mascaro AL, Silvestri L, Müllenbroich MC, et al. A versatile clearing agent for multi-modal brain imaging. Sci Rep. 2015;5: 9808.

[23] Lee E, Choi J, Jo Y, Kim JY, Jang YJ, Lee HM, et al. ACT-PRESTO: Rapid and consistent tissue clearing and labeling method for 3-dimensional (3D) imaging. Sci Rep 2016;6: 18631.

[24] Susaki EA, Tainaka K, Perrin D, Kishino F, Tawara T, Watanabe TM. Whole-brain imaging with single-cell resolution using chemical cocktails and computational analysis. Cell. 2014;157: 726–739.

[25] Tainaka K, Kubota SI, Suyama TQ, Susaki EA, Perrin D, Ukai-Tadenuma M. Whole-body imaging with single-cell resolution by tissue decolorization. Cell. 2014;159: 911–924.

[26] Kubota SI, Takahashi K, Nishida J, Morishita Y, Ehata S, Tainaka K, et al. Whole-Body Profiling of Cancer Metastasis with Single-Cell Resolution. Cell Rep. 2017;20: 236–250.

[27] Matryba P, Bozycki L, Pawłowska M, Kaczmarek L, Stefaniuk M. Optimized perfusion-based CUBIC protocol for the efficient whole-body clearing and imaging of rat organs. J Biophotonics. 2018;11: e201700248.

[28] Hama H, Hioki H, Namiki K, Hoshida T, Kurokawa H, Ishidate F, et al. ScaleS: an optical clearing palette for biological imaging. Nat Neurosci. 2015;18: 1518–1529.

[29] Tainaka K, Murakami TC, Susaki EA, Shimizu C, Saito R, Takahashi K. Chemical Landscape for Tissue Clearing Based on Hydrophilic Reagents. Cell Rep. 2018;24: 2196–2210.

[30] Susaki EA, Ueda HR. Whole-body and whole-organ clearing and imaging techniques with single-cell resolution: Toward organism-level systems biology in mammals. Cell Chem Biol. 2016;23: 137–157.

[31] Bangham A. Properties and uses of lipid vesicles: an overview. Ann N Y Acad Sci. 1978;308: 2–7.

[32] Enoki S, Saeki K, Maki K, Kuwajima K. Acid denaturation and refolding of green fluorescent protein. Biochemistry. 2004;43: 14238–14248.

[33] Schwarz MK, Scherbarth A, Sprengel R, Engelhardt J, Theer P, Giese G. Fluorescent-protein stabilization and high-resolution imaging of cleared, intact mouse brains. PLoS One. 2015;10:e0124650.

